# Tumor suppressor p53 controls thymic NKT17 development

**DOI:** 10.1101/2024.08.21.608967

**Authors:** Sofia Celli, Masashi Watanabe, Richard J. Hodes

## Abstract

The tumor suppressor p53 antagonizes tumorigenesis, notably including the suppression of T cell lymphomas while its role on physiological T cell biology including thymic T cell development has not been fully understood. Invariant natural killer T (iNKT) cells develop in the thymus as innate-like αβ-T cells which consist of NKT1, NKT2 and NKT17 subsets. We found that the tumor suppressor p53 regulates specifically thymic NKT17 development. p53 is highly expressed in NKT17 relative to other T cell populations. Loss of p53 in the T cell lineage resulted in increased thymic NKT17 cell number with retention of lineage specific cytokine production, while development of NKT1, NKT2 and conventional T cells was not affected. Of interest, γH2AX expression was higher in NKT17 than NKT1 and NKT2 at steady state, and it was further increased in p53-deficient NKT17, suggesting that NKT17 development involves selectively greater DNA damage or genomic instability and that p53 expression might be in response to these damage signals. Taken together, our results indicated that the tumor suppressor p53 is active in selectively controlling thymic NKT17 development, with absence of p53 leading to an increase in thymic NKT17 cells expressing high levels of DNA damage response.

## INTRODUCTION

Tumor suppressor p53 regulates cell proliferation and cell death in tumorigenesis and is the most frequent mutation observed in human cancers (Kastenhuber and Lowe, 2017; Oren and Prives, 2024). p53 null mice spontaneously develop tumors including lethal thymic lymphoma by three months of age (Nacht et al., 1996). Although extensive cell proliferation and cell death occur during thymic T cell development, there was no obvious T cell developmental alteration observed in p53 deficient mice (Lowe et al., 1993; Nacht et al., 1996) although a role for p53 among all T cell subsets has not been fully studied.

Invariant natural killer T cells (iNKT) are αβ-T cells expressing semi-invariant TCR and possessing innate-like functional phenotypes (Bendelac et al., 1997). The αβ-TCRs of iNKT are characterized by expression of Vα14-Jα18 in mouse and Vα24-Jα18 in human, with biased TCRβ usage (Taniguchi et al., 2003). iNKT cells develop in the thymus as three distinct subsets: NKT1, which are T-bet^+^ and produce IFNγ; NKT2, which are GATA3^+^ PLZF^high^ and produce IL-4; and NKT17, which are RORγt^+^ and produce IL-17 (Hogquist and Georgiev, 2020). The relative proportions of thymic NKT1, NKT2 and NKT17 cells differ across mouse strains and across tissues (Lee et al., 2015), and several models have been proposed to explain the differentiation of distinct NKT subsets (Hogquist and Georgiev, 2020; Krovi et al., 2022). While NKT17 is the smallest population among NKT subsets in the thymus (Lee et al., 2015), the regulatory mechanisms that control the size of the thymic NKT17 population have not been completely understood (Liman and Park, 2023; Tsagaratou, 2019). In the present study, we demonstrated that the tumor suppressor p53 regulates development of thymic NKT17 cells expressing high levels of DNA damage response.

## RESULTS

### p53 expression in thymic T cells

We assessed the role of p53 in the development of thymic T cell subpopulations. We first analyzed p53 protein expression by flow cytometry (Watanabe et al., 2014) in thymic T cell sub-populations including conventional T cell lineages (DN, DP, CD4SP, CD8SP) as well as non-conventional T cell lineage iNKT cells (**Figure 1A and 1B**). Thymic iNKT cells were detected by PBS57-CD1d tetramer binding, and iNKT subsets (NKT1, NKT2 and NKT17) were identified by expression pattern of PLZF and RORγt transcription factors (Watanabe et al., 2022) (**Figure 1B**). At steady state, NKT17 cells showed the highest level of p53 protein expression compared to NKT1, NKT2 and CD4/CD8 double positive (DP) thymocytes (**Figure 1C**). It is known that p53 is upregulated in thymocytes by irradiation and mediates irradiation-induced thymocyte apoptosis (Lowe et al., 1993). Indeed, following irradiation, p53 upregulation was observed in NKT1, NKT2, NKT17 and DP cells, with irradiated-NKT17 retaining the highest level of p53 protein expression (**Figure 1C**). These results demonstrated that p53 protein expression is selectively upregulated in NKT17 at steady state. Of interest, we also observed that γH2AX expression, a marker of genomic instability which may trigger p53 activation, was also higher in NKT17 cells (**Figure 1D**). These results suggested that p53 might have a differential role in thymic NKT17 cell development, potentially reflecting a response to a high level of DNA damage in this lineage.

**Figure 1.**
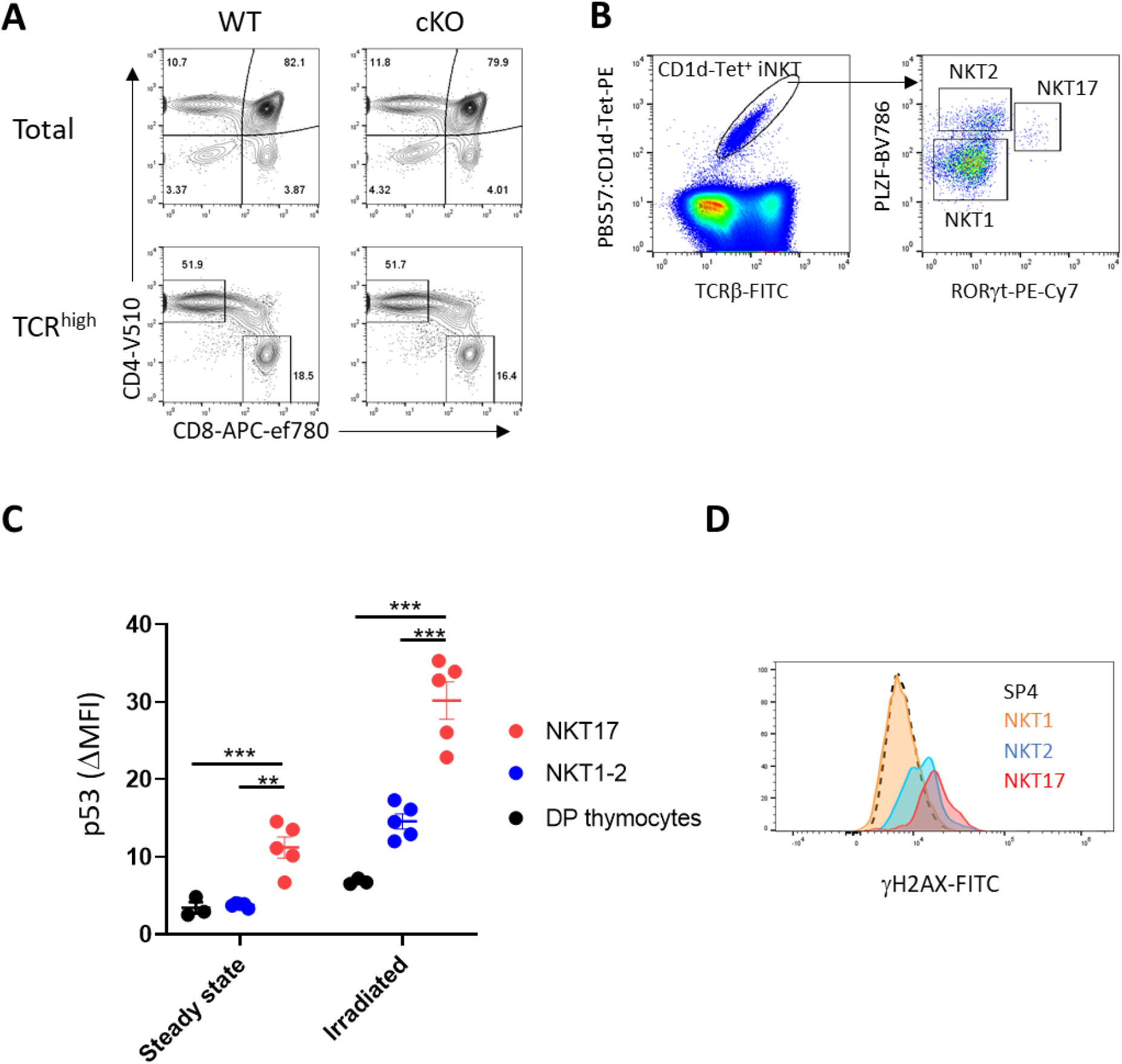
NKT17 expresses higher level of p53 protein than other T cell subsets. (A) Conventional T cells develop normally in the absence of p53. Upper panel, CD4 and CD8 profile of total thymocytes. Lower panel, CD4 and CD8 profile of TCRβ ^high^ matured thymocytes. (B) Representative gating strategy for thymic NKT1 , NKT2 and NKT17 subsets. (C) p53 protein expression levels were analyzed by flow cytometry at steady state and after irradiation. For irradiation condition, cells were irradiated (1,000 rad) and incubated for 4 hours at 37°C. Data is pooled result of two independent experiments. (D) γH2AX expression in thymic iNKT subset was analyzed by flow cytometry. Data are mean ± SEM. Statistical differences between groups were analyzed with One-way ANOVA followed by multiple comparison. * * *p* < 0.005, * * * *p* < 0.0001.

### Loss of p53 results in increase of thymic NKT17

To directly test the role of the tumor suppressor p53 in thymic T cell development, the *Trp53* gene was specifically deleted from T cell lineage by crossing B6 p53^flox^ and CD4-Cre strains (hereafter described as p53 cKO) in which the early onset of thymic lymphoma development and lethality observed in p53 null mice is prevented (Kawashima et al., 2013). In the p53 cKO mice, there was no detectable change in the CD4 / CD8 profile of thymic T cells and no developmental defect in TCRβ^high^ CD4 or CD8 single positive mature thymocytes (**Figure 1A**), consistent with previous reports (Lowe et al., 1993; Nacht et al., 1996). While the total iNKT cell population size did not change in the absence of p53 (**Figure 2, bottom left**), NKT17 (PLZF^int^, RORγt^+^) was significantly increased (**Figure 2, bottom right**). In a preliminary analysis, we observed that the expanded NKT17 cells in p53 cKO mice exhibited IL-17 production comparable to p53 WT NKT17 upon *in vitro* stimulation (data not shown). Considering the transcriptional similarity between iNKT and MAIT cell subsets (Godfrey et al., 2019), MAIT cell subsets (MAIT1 and MAIT17) were also analyzed in p53 cKO mice with 5-OP-RU/MR1 tetramer; however, there were no differences in MAIT1 (T-bet^+^) and MAIT17 (RORγt^+^) between p53 cKO and WT control thymus (data not shown). These results indicated that loss of p53 during T cell development specifically affects thymic NKT17 cells.

**Figure 2.**
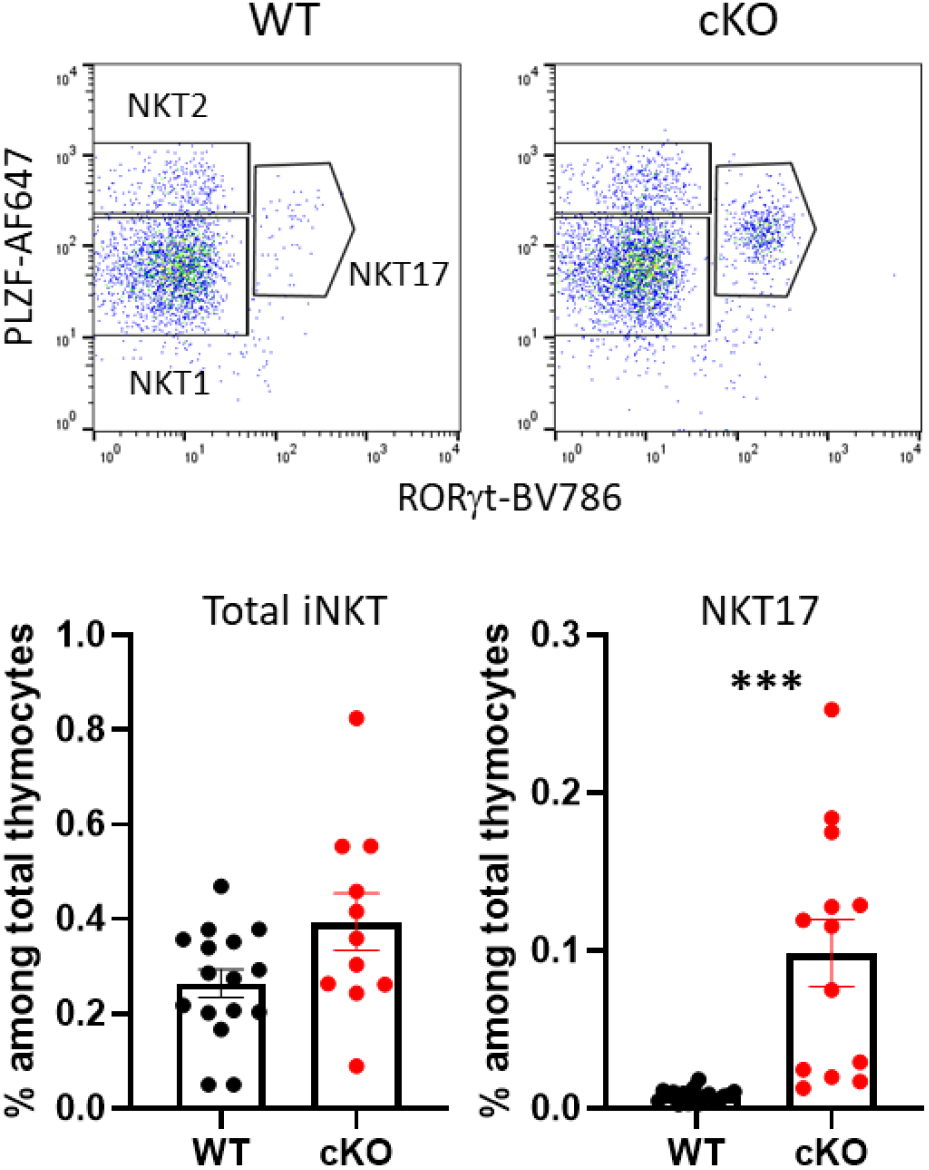
Loss of p53 results in increase of thymic NKT17 cells. Upper panel, representative FACS plot of iNKT subsets of p53 cKO strain. Lower panel, pooled data of ten independent experiments for frequency of total iNKT and NKT17 cells among total thymocytes. Data are mean ± SEM. Statistical differences between groups were analysed with two-tailed student t-test. * * * *p* < 0.0001.

### p53 deficiency impacts thymic NKT17 but not peripheral NKT17

To understand whether p53 deficiency selectively affects NKT17 cells residing in the thymus or if it also impacts peripheral NKT17 populations, NKT17 cells in peripheral organs were analyzed. It has been previously reported that distinct iNKT subsets are preferentially enriched in different peripheral tissues (Lee et al., 2015); for instance, NKT17 cells are most abundant in lymph nodes and lung in the B6 strain. In lymph nodes and lung, there was no significant difference in the NKT17 population between p53 intact and cKO mice (**Figure 3A and 3B**) indicating a unique role for p53 in regulating the population size of thymic NKT17.

**Figure 3.**
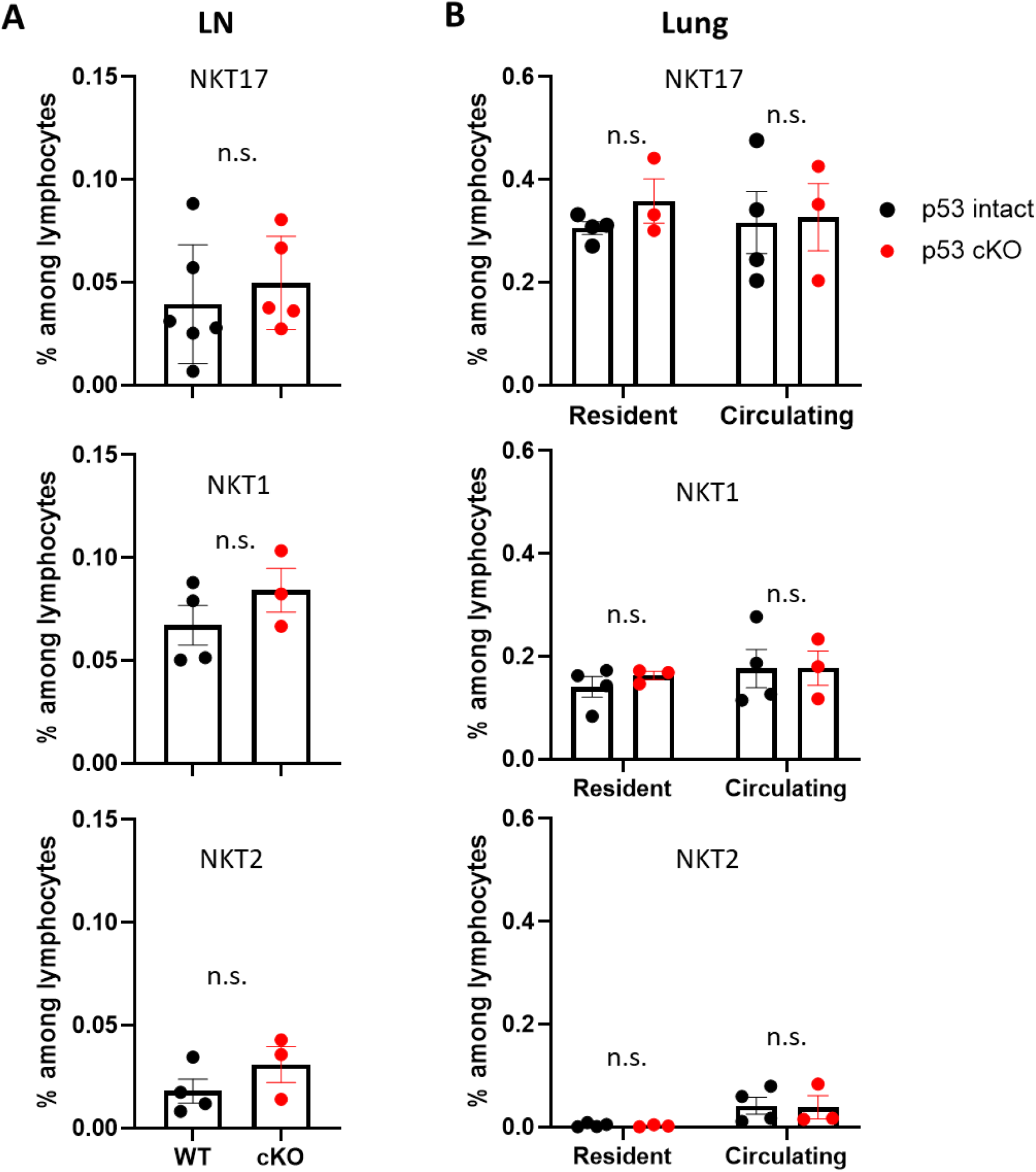
p53 deficiency does not affect NKT17 population in peripheral tissues. (A) NKT17 cells in peripheral inguinal lymph nodes. Data is pooled result of three independent experiments. (B) NKT17 cells in lung tissue. Mice were i.v. injected anti-CD45-AF488 Ab to label circulating lymphocyte in blood. Five minutes after injection, lung was dissected, and lymphocytes were isolated from lung. Circulating NKT17 was defined as CD45-AF488 positive while resident NKT17 was defined as CD45-AF488 negative cells. Data is pooled result of three independent experiments. Data are mean ± SEM. Statistical differences between groups were analyzed with student t-test. n.s., not significant.

### Mechanistic basis for increase of thymic NKT17 in the absence of p53

Next, the mechanism by which p53-deficiency led to significantly increased thymic NKT17 was investigated. It was reported that IL-7 receptor signaling is important for NKT17 development by inducing their expansion and supporting homeostasis (Webster et al., 2014). We previously reported that p53 negatively controls cytokine-induced bystander T cell proliferation in certain situations (Watanabe et al., 2014). To test whether NKT17 proliferation was enhanced in the absence of p53, Ki67 expression was analyzed as an indicator of proliferating cells. Although thymic NKT2 and NKT17 but not NKT1 showed a Ki67^+^ proliferating phenotype, there was no difference in frequency of Ki67^+^ cells between p53-deficient and p53-intact NKT17 and NKT2 (**Figure 4A**). This result indicated that increased NKT17 in the p53 cKO thymus was not directly related to enhanced proliferation. As a tumor suppressor, p53 senses abnormal cellular stress, such as oncogene activation and/or DNA damages, and subsequently induces cell cycle arrest and/or cell death to prevent tumor development (Kruse and Gu, 2009). We hypothesized that DNA damage response might be involved and that p53 might have a role in eliminating a proportion of NKT17 cells during thymic development. Consistent with this possibility, thymic NKT17 expressed the highest level of γH2AX, a marker of DNA damage / genomic instability, among iNKT subsets (**Figure 1D and 4B**). Of interest, γH2AX level was further increased in p53-deficient NKT17 compared to p53-intact NKT17 (**Figure 4B**). These results suggested that NKT17 cells are more prone to DNA damage / genomic instability marked by γH2AX during thymic development. This feature of NKT17 might lead to activation of p53-dependent effector pathway(s) to eliminate a portion of thymic NKT17 cells, controlling the size of this population.

**Figure 4.**
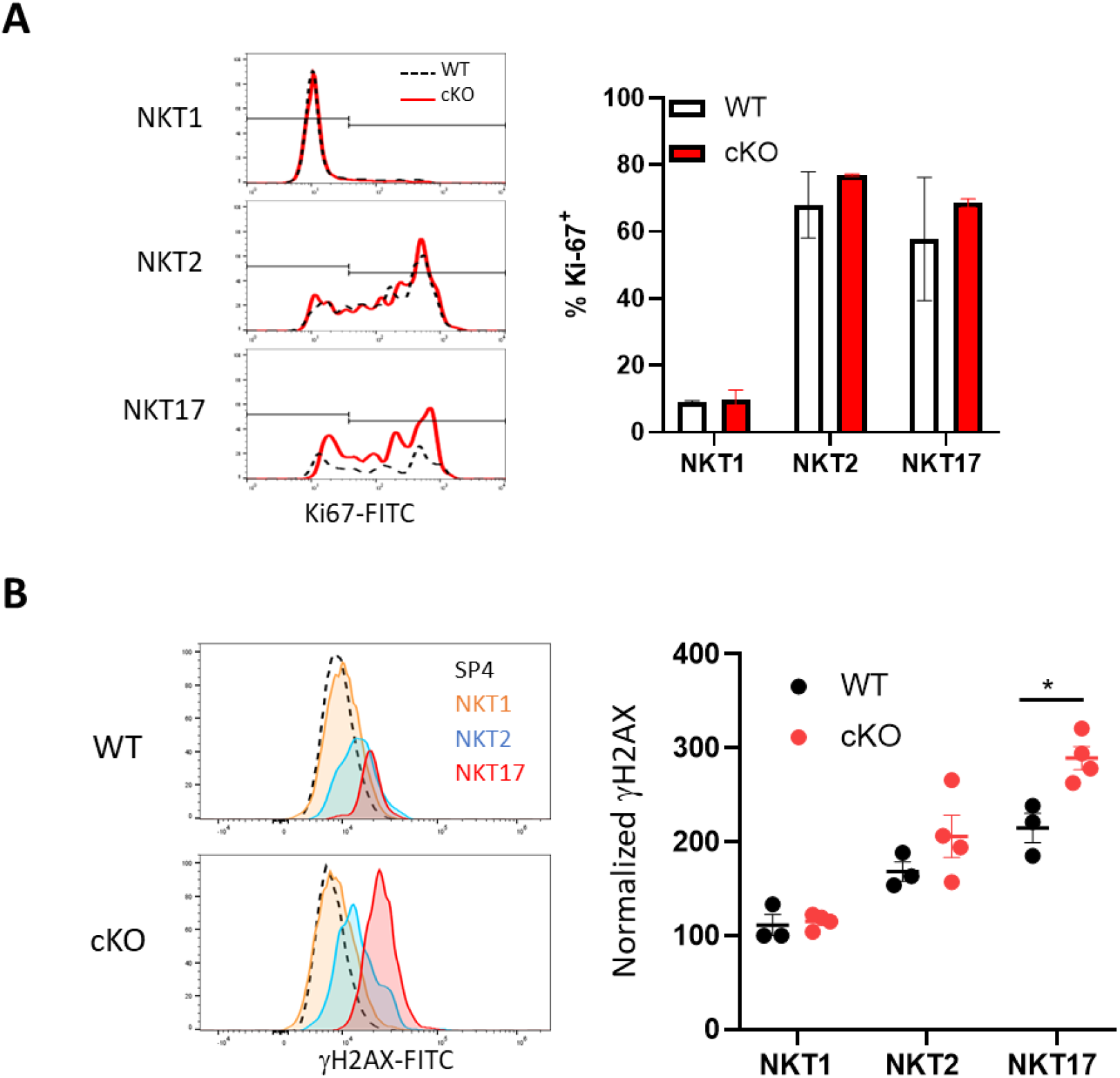
γH2AX is higher in NKT17 and further accumulated in the absence of p53. (A) Ki67 expression in thymic iNKT subset was analyzed by flow cytometry. Data is pooled result of two independent experiments. WT, n=2; cKO, n=2. (B) γH2AX expression in thymic iNKT subset was analyzed by flow cytometry. Data is pooled result of three independent experiments. SP4; CD4 single positive thymocytes. WT, n=3; cKO, n=4. Data are mean ± SEM. Statistical differences were analyzed with student t-test. * *p* < 0.05.

## DISCUSSION

In the present study, we showed that the loss of tumor suppressor p53 in T cell-lineage resulted in significantly increased thymic NKT17 cell number. p53 regulates cell proliferation and cell death in tumorigenesis and is the most frequently mutated gene in human cancers (Kastenhuber and Lowe, 2017; Oren and Prives, 2024). p53 null mice spontaneously develop tumors including lethal thymic lymphoma within a few months of age (Nacht et al., 1996). Although extensive cell proliferation and cell death occur during thymic T cell development, previous studies reported no alteration in T cell development in p53 deficient mice (Lowe et al., 1993; Nacht et al., 1996). To our knowledge, the study presented here is the first report of p53 regulation of a specific thymic T cell subset, evidenced by the large selective expansion of thymic NKT17 in p53 cKO mice.

At steady state, p53 protein level is strictly regulated by E3 ubiquitin ligase MDM2 through a proteasome degradation pathway (Kruse and Gu, 2009). Upon genomic instability and/or oncogene activation, MDM2 regulation of p53 is interrupted, resulting in upregulation and activation of p53 followed by activation of downstream p53 effector pathways (Kruse and Gu, 2009). We found that the level of p53 protein in thymic NKT17 was the highest among thymic T cell subsets analyzed, both *ex vivo* and after irradiation. These results suggested the presence of a higher level of cellular stress leading to upregulation of p53 protein in thymic NKT17. We did not find evidence of altered cellular proliferation of p53-deficient thymic NKT17 cells. γH2AX expression, a marker of genomic instability, was higher in p53-intact NKT17 compared to NKT1 and NKT2 and was further accumulated in p53-deficient NKT17 cells. This suggested that thymic NKT17 cells are more prone to genomic instability during thymic development and that p53 actively controls NKT17 population size.

In our preliminary observation, p53 cKO thymic NKT17 cells exhibited a consistent lineage specific cytokine production pattern upon *in vitro* stimulation, indicating that p53 did not affect lineage specific function. While peripheral functions of NKT17 have been reported (Tsagaratou, 2019), the precise role of thymic NKT17 and their production of cytokines including IL-17 in thymus has not been well understood. In addition to increasing population size, whether p53-deficiency affects functional roles of thymic NKT17, if any, remains to be addressed.

It was reported that murine peripheral T cell lymphomas (PTCLs) develop in p53 KO mice and originate from a CD1d-restricted NKT cell lineage with capability to produce IL-17 (Bachy et al., 2016). Although it was speculated that the PTCL development was a post-thymic event (Bachy et al., 2016), the significant increase of thymic NKT17 number in the absence of p53 reported in this study might also contribute to increase to the risk of PTCL development.

Taken together, this current study demonstrates a novel role for the tumor suppresser p53 in controlling the population size of thymic NKT17 cells.

## MATERIALS AND METHODS

### Mice

C57BL/6 (B6) mice were purchased from Charles River. B6 p53 KO (B6.129S2-Trp53tm1Tyj/J, strain#:002101), B6 p53-flox (B6.129P2-Trp53tm1Brn/J, strain#: 008462), CD4-Cre (B6.Cg-Tg(Cd4-cre)1Cwi/BfluJ, Strain #:022071) and IL-17A-EGFP (C57BL/6-Il17atm1Bcgen/J, Strain #:018472) mice were purchased from Jaxson Laboratory. Mice were maintained in accordance with National Institutes of Health guidelines. All animal experiments were approved by the National Cancer Institute Animal Care and Use Committees.

### Lung lymphocyte isolation

Mice were injected intravenously with 2.5 μg of fluorochrome-labeled anti-CD45.2 or anti-CD45 antibody, and after 3 min lungs were harvested (Sakai et al., 2016). Briefly, lungs were minced with scissors and then enzymatically digested for 45 min at 37°C in RPMI-1640 medium supplemented with 1 mg/ml Collagenase D (Roche-Diagnostics, Indianapolis, IN), 1 mg/ml hyaluronidase, 50 U/ml DNase I and 1 mM aminoguanidine (all from Sigma-Aldrich, St. Louis, MO). Digested lung was dispersed by passage through a 100 μm pore size cell-strainer and lung lymphocytes were enriched by 37% Percoll density centrifugation.

### *In vitro* stimulation

Thymic iNKT cells were stimulated as previously reported (Watanabe et al., 2022). Briefly, 2 x 10^6^ total thymocytes were stimulated with PMA (5 ng/mL) and Ionomycin (1000 ng/mL) in a 24-well plate for 4 hours at 37 °C. Golgi Stop (0.67 μl/mL) and Golgi Plug (1 μl/mL) (eBioscience/Thermo Fisher) were added after 1 hour of incubation.

### Flow Cytometry

Cells were washed with FACS buffer (HBSS containing 0.2% BSA and 0.05% Azide), treated with anti-FcR (2.4G2), and then stained with the following Abs. Anti-CD4 (RM4-5)-BV510 (Need clone name or described it in Star Protocol), anti-B220 (RA3-6B2)-AF700, anti-CD44 (IM7)-FITC, anti-TCRβ (H57-597)-PE/Cy7, anti-RORγt (Q31-378)-BV786, anti-T-bet (O4-46)-BV421, anti-PLZF (R17-809)-AF647, anti-CCR7 (4B12)-BV786, anti-IL-17A (TC11-18H10)-FITC, anti-IFNγ (XMG1.2)-FITC, anti-IL-4 (11B11)-BV421 were purchased from BD Biosciences. Anti-NK1.1 (PK136)-APC/eFluor780, anti-CD8a (53-6-7)-APC/eFluor780, anti-B7.1 (16-10A1)-APC, anti-B7.2 (GL1)-APC, anti-CD28 (37.51)-APC, anti-Egr2 (erongr2)-APC were purchased from eBioscience/Thermo Fisher. Anti-CD24 (30-F1)-AF594 and anti-CD69 (H1.2F3)-BV421 were purchased from BioLegend. For detection of iNKT cells, PE-CD1d tetramer loaded with PBS57 (α-galactosyl ceramide analogue), obtained from the NIH Tetramer Core Facility, was used. For intracellular staining, Foxp3 staining kit (eBioscience/Thermo Fisher) was used for fixation and permeabilization according to manufacturer’s instructions and then stained with Abs for 30 min at 4 °C. Data were collected with a FACS Fortessa (BD Biosciences) flow cytometer and analyzed with FlowJo (version 10.8) software (Tree Star).

### Statistics

For statistical analysis, Prism GraphPad (version 8.3.4) software was used. Student’s t test with two-tailed distribution was performed for statistical analyses with a single comparison. For multiple comparisons, statistical analysis was performed with one-way ANOVA followed by Dunnett’s multiple comparison. *p* values < 0.05 were considered statistically significant.

## ACKNOWLEDGEMENTS

We thank NCI Frederick animal facility for care and maintenance of animal.

## AUTHOR CONTRIBUTIONS

S.C. and M.W. designed and performed experiments, analyzed results, and wrote the manuscript. R.J.H supervised the study and writing of the manuscript.

## REFERENCES

Bachy, E., M. Urb, S. Chandra, R. Robinot, G. Bricard, S. de Bernard, A. Traverse-Glehen, S. Gazzo, O. Blond, A. Khurana, L. Baseggio, T. Heavican, M. Ffrench, G. Crispatzu, P. Mondiere, A. Schrader, M. Taillardet, O. Thaunat, N. Martin, S. Dalle, M. Le Garff-Tavernier, G. Salles, J. Lachuer, O. Hermine, V. Asnafi, M. Roussel, T. Lamy, M. Herling, J. Iqbal, L. Buffat, P.N. Marche, P. Gaulard, M. Kronenberg, T. Defrance, and L. Genestier. 2016. CD1d-restricted peripheral T cell lymphoma in mice and humans. J Exp Med 213:841–857.

Bendelac, A., M.N. Rivera, S.H. Park, and J.H. Roark. 1997. Mouse CD1-specific NK1 T cells: development, specificity, and function. Annu Rev Immunol 15:535–562.

Dai, H., A. Rahman, A. Saxena, A.K. Jaiswal, A. Mohamood, L. Ramirez, S. Noel, H. Rabb, C. Jie, and A.R. Hamad. 2015. Syndecan-1 identifies and controls the frequency of IL-17-producing naive natural killer T (NKT17) cells in mice. Eur J Immunol 45:3045–3051.

Godfrey, D.I., H.F. Koay, J. McCluskey, and N.A. Gherardin. 2019. The biology and functional importance of MAIT cells. Nat Immunol 20:1110–1128.

Hogquist, K., and H. Georgiev. 2020. Recent advances in iNKT cell development. F1000Res 9:

Kastenhuber, E.R., and S.W. Lowe. 2017. Putting p53 in Context. Cell 170:1062–1078.

Kawashima, H., H. Takatori, K. Suzuki, A. Iwata, M. Yokota, A. Suto, T. Minamino, K. Hirose, and H. Nakajima. 2013. Tumor suppressor p53 inhibits systemic autoimmune diseases by inducing regulatory T cells. J Immunol 191:3614–3623.

Krovi, S.H., L. Loh, A. Spengler, T. Brunetti, and L. Gapin. 2022. Current insights in mouse iNKT and MAIT cell development using single cell transcriptomics data. Semin Immunol 60:101658.

Kruse, J.P., and W. Gu. 2009. Modes of p53 regulation. Cell 137:609–622.

Lee, Y.J., H. Wang, G.J. Starrett, V. Phuong, S.C. Jameson, and K.A. Hogquist. 2015. Tissue-Specific Distribution of iNKT Cells Impacts Their Cytokine Response. Immunity 43:566–578.

Liman, N., and J.H. Park. 2023. Markers and makers of NKT17 cells. Exp Mol Med 55:1090–1098.

Lowe, S.W., E.M. Schmitt, S.W. Smith, B.A. Osborne, and T. Jacks. 1993. p53 is required for radiation-induced apoptosis in mouse thymocytes. Nature 362:847–849.

Luo, S., J. Kwon, A. Crossman, P.W. Park, and J.H. Park. 2021. CD138 expression is a molecular signature but not a developmental requirement for RORgammat+ NKT17 cells. JCI Insight 6:

Nacht, M., A. Strasser, Y.R. Chan, A.W. Harris, M. Schlissel, R.T. Bronson, and T. Jacks. 1996. Mutations in the p53 and SCID genes cooperate in tumorigenesis. Genes Dev 10:2055–2066.

Oren, M., and C. Prives. 2024. p53: A tale of complexity and context. Cell 187:1569–1573.

Sakai, S., K.D. Kauffman, M.A. Sallin, A.H. Sharpe, H.A. Young, V.V. Ganusov, and D.L. Barber. 2016. CD4 T Cell-Derived IFN-gamma Plays a Minimal Role in Control of Pulmonary Mycobacterium tuberculosis Infection and Must Be Actively Repressed by PD-1 to Prevent Lethal Disease. PLoS Pathog 12:e1005667.

Taniguchi, M., M. Harada, S. Kojo, T. Nakayama, and H. Wakao. 2003. The regulatory role of Valpha14 NKT cells in innate and acquired immune response. Annu Rev Immunol 21:483–513.

Tsagaratou, A. 2019. Unveiling the regulation of NKT17 cell differentiation and function. Mol Immunol 105:55–61.

Watanabe, M., S. Celli, F.A. Alkhaleel, and R.J. Hodes. 2022. Antigen-presenting T cells provide critical B7 co-stimulation for thymic iNKT cell development via CD28-dependent trogocytosis. Cell Rep 41:111731.

Watanabe, M., K.D. Moon, M.S. Vacchio, K.S. Hathcock, and R.J. Hodes. 2014. Downmodulation of tumor suppressor p53 by T cell receptor signaling is critical for antigen-specific CD4(+) T cell responses. Immunity 40:681–691.

Webster, K.E., H.O. Kim, K. Kyparissoudis, T.M. Corpuz, G.V. Pinget, A.P. Uldrich, R. Brink, G.T. Belz, J.H. Cho, D.I. Godfrey, and J. Sprent. 2014. IL-17-producing NKT cells depend exclusively on IL-7 for homeostasis and survival. Mucosal Immunol 7:1058–1067.

